# Re-assembly, quality evaluation, and annotation of 678 microbial eukaryotic reference transcriptomes

**DOI:** 10.1101/323576

**Authors:** Lisa K. Johnson, Harriet Alexander, C. Titus Brown

**Affiliations:** Department of Population Health & Reproduction, School of Veterinary Medicine, University of California Davis; Molecular, Cellular, and Integrative Physiology Graduate Group, University of California Davis; Genome Center, University of California Davis

**Keywords:** marine microbial eukaryote, transcriptome assembly, automated pipeline, re-analysis

## Abstract

**Background:** *De novo* transcriptome assemblies are required prior to analyzing RNAseq data from a species without an existing reference genome or transcriptome. Despite the prevalence of transcriptomic studies, the effects of using different workflows, or “pipelines”, on the resulting assemblies are poorly understood. Here, a pipeline was programmatically automated and used to assemble and annotate raw transcriptomic short read data collected by the Marine Microbial Eukaryotic Transcriptome Sequencing Project (MMETSP). The resulting transcriptome assemblies were evaluated and compared against assemblies that were previously generated with a different pipeline developed by the National Center for Genome Research (NCGR).

**Results:** New transcriptome assemblies contained the majority of previous contigs as well as new content. On average, 7.8% of the annotated contigs in the new assemblies were novel gene names not found in the previous assemblies. Taxonomic trends were observed in the assembly metrics, with assemblies from the Dinoflagellata and Ciliophora phyla showing a higher percentage of open reading frames and number of contigs than transcriptomes from other phyla.

**Conclusions:** Given current bioinformatics approaches, there is no single ‘best’ reference transcriptome for a particular set of raw data. As the optimum transcriptome is a moving target, improving (or not) with new tools and approaches, automated and programmable pipelines are invaluable for managing the computationally-intensive tasks required for re-processing large sets of samples with revised pipelines and ensuring a common evaluation workflow is applied to all samples. Thus, re-assembling existing data with new tools using automated and programmable pipelines may yield more accurate identification of taxon-specific trends across samples in addition to novel and useful products for the community.

**Key Points:**

- Re-assembly with new tools can yield new results
- Automated and programmable pipelines can be used to process arbitrarily many samples.
- Analyzing many samples using a common pipeline identifies taxon-specific trends.

## Introduction

The analysis of gene expression from high-throughput nucleic acid sequence data relies on the presence of a high quality reference genome or transcriptome. When there is no reference genome or transcriptome for an organism of interest, raw RNA sequence data (RNAseq) must be assembled *de novo* into a transcriptome [1]. This type of analysis is ubiquitous across many fields, including: evolutionary developmental biology [2], cancer biology [3], agriculture [4, 5], ecological physiology [6, 7], and biological oceanography [8]. In recent years, substantial investments have been made in data generation, primary data analysis, and development of downstream applications, such as biomarkers and diagnostic tools [9, 10, 11, 12, 13, 14, 15, 16]

Methods for *de novo* RNAseq assembly of the most common short read Illumina sequencing data continue to evolve rapidly, especially for non-model species [17]. At this time, there are several major *de novo* transcriptome assembly software tools available to choose from, including Trinity [18], SOAPdenovo-Trans [19], Trans-ABySS [20], Oases [21], SPAdes [22], IDBA-tran [23], and Shannon [24]. The availability of these options stems from continued research into the unique computational challenges associated with transcriptome assembly of short read Illumina RNAseq data, including large memory requirements, alternative splicing and allelic variants [18, 25],

The continuous development of new tools and workflows for RNAseq analysis combined with the vast amount of publicly available RNAseq data [26] raises the opportunity to re-analyze existing data with new tools. This, however, is rarely done systematically. To evaluate the performance impact of new tools on old data, we developed and applied a programmatically automated *de novo* transcriptome assembly workflow that is modularized and extensible based on the Eel Pond Protocol [27]. This workflow incorporates Trimmomatic [28], digital normalization with khmer software [29, 30], and the Trinity *de novo* transcriptome assembler [18].

To evaluate this pipeline, we re-analyzed RNAseq data from 678 samples generated as part of the Marine Microbial Eukaryotic Transcriptome Sequencing Project (MMETSP) [31]. The MMETSP data set was generated to broaden the diversity of sequenced marine protists to enhance our understanding of their evolution and roles in marine ecosystems and biogeochemical cycles [31, 32]. With data from species spanning more than 40 eukaryotic phyla, the MMETSP provides one of the largest publicly-available collections of RNAseq data from a diversity of species. Moreover, the MMETSP used a standardized library preparation procedure and all of the samples were sequenced at the same facility, making this data set unusually comparable.

Reference transcriptomes for the MMETSP were originally assembled by the National Center for Genome Research (NCGR) with a pipeline which used the Trans-ABySS software program to assemble the short reads [31]. The transcriptomes generated from the NCGR pipeline have already facilitated discoveries in the evolutionary history of ecologically significant genes [33, 34], differential gene expression under shifting environmental conditions [8, 35], inter-group transcriptomic comparisons [36], unique transcriptional features [37, 38, 39], and meta-transcriptomic studies [34, 35, 36]

In re-assembling the MMETSP data, we sought to compare and improve the original MMETSP reference transcriptome and to create a platform which facilitates automated re-assembly and evaluation. Here, we show that our re-assemblies had better evaluation metrics and contained most of the NCGR contigs as well as adding new content.

## Methods

### Programmatically Automated Pipeline

An automated pipeline was developed to execute the steps of the Eel Pond mRNAseq Protocol [27], a lightweight protocol for assembling short Illumina RNAseq reads that uses the Trinity *de novo* transcriptome assembler. This protocol generates *de novo* transcriptome assemblies of acceptable quality [40]. The pipeline was used to assemble all of the data from the MMETSP (Figure 1). The code and instructions for running the pipeline are available at https://doi.org/10.5281/zenodo.740440 [41].

**Figure 1.**
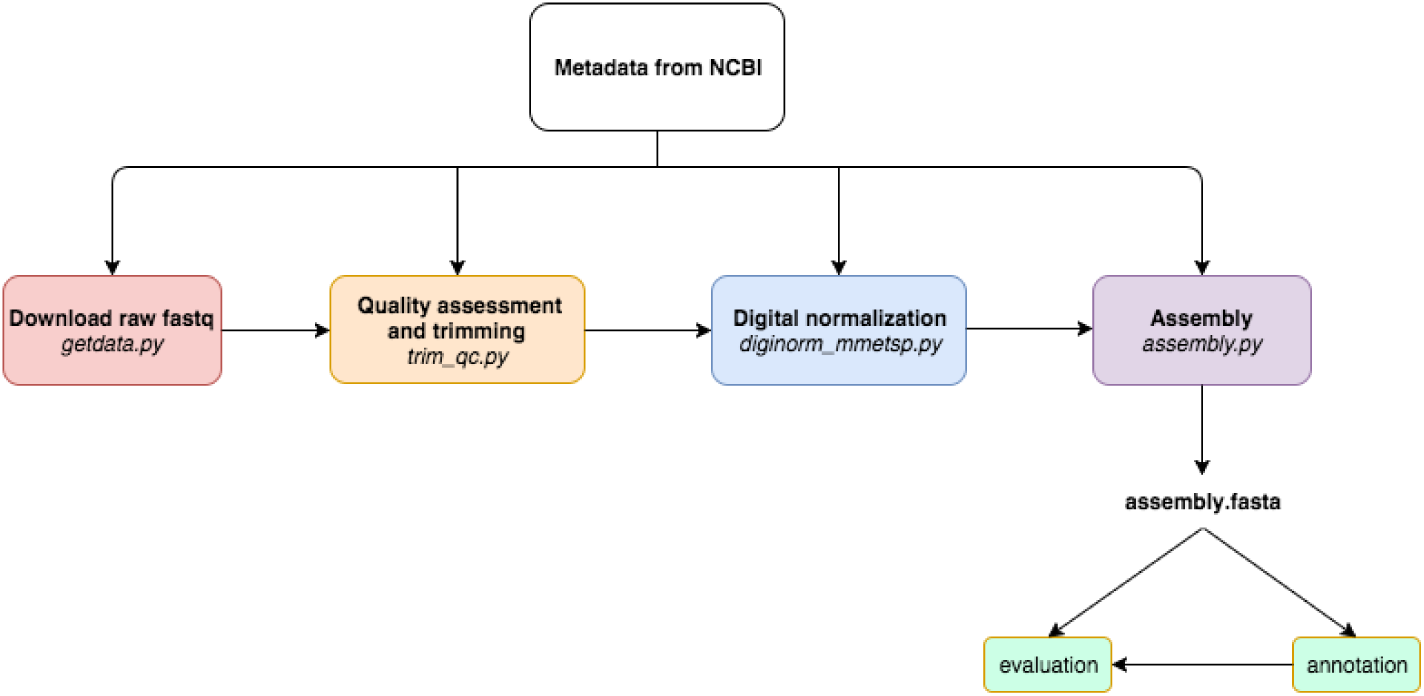
A programmatically automated *de novo* transcriptome assembly pipeline was developed for this study. Metadata in the SraRunInfo.csv file downloaded from NCBI was used as input for each step of the pipeline to indicate which samples were processed. The steps of the pipeline are as follows: download raw fastq data with the fastq-dump script in the SRA Toolkit, perform quality assessment with FastQC and trim residual Illumina adapters and low quality bases (Q<2) with Trimmomatic, do digital normalization with khmer version 2.0, and perform *de novo* transcriptome assembly with Trinity. If a process was terminated, the automated nature of this pipeline allowed for the last process to be run again without starting the pipeline over. In the future, if a new sample is added, the pipeline can be run from beginning to end with just new samples, without having to repeat the processing of all samples in the dataset as one batch. If a new tool becomes available, for example a new assembler, it can be substituted in lieu of the original tool used by this pipeline.

The steps of the pipeline applied to the MMETSP are as follows:

### 1. Download the raw data

Raw RNA-seq data sets were obtained from the National Center for Biotechnology Information (NCBI) Sequence Read Archive (SRA) from BioProject PRJNA231566. Data were paired-end (PE) Illumina reads with lengths of 50 bases for each read. A metadata (SraRunInfo.csv) file obtained from the SRA web interface was used to provide a list of samples to the get_data.py pipeline script, which was then used to download and extract fastq files from 719 records. The script uses the fastq-dump program from the SRA Toolkit to extract the SRA-formatted fastq files (version 2.5.4) [42]. There were 18 MMETSP samples with more than one SRA record (MMETSP0693, MMETSP1019, MMETSP0923, MMETSP0008, MMETSP1002, MMETSP1325, MMETSP1018, MMETSP1346, MMETSP0088, MMETSP0092, MMETSP0717, MMETSP0223, MMETSP0115, MMETSP0196, MMETSP0197, MMETSP0398, MMETSP0399, MMETSP0922). In these cases, reads from multiple SRA records were concatenated together per sample. Taking these redundancies into consideration, there were a total of 678 re-assemblies generated from the 719 records in PRJNA231566 (Supplemental Notebook 1 [43]). Assembly evaluation metrics were not calculated for MMETSP samples with more than one SRA record because these assemblies were different than the others, containing multiple samples, and thus not as comparable.

Initial transcriptomes that were assembled by the National Center for Genome Resources (NCGR), using methods and data described in the original publication [31], were downloaded from the iMicrobe repository to compare with our re-assemblies (ftp://ftp.imicrobe.us/projects/104/). There were two versions of each assembly, ‘nt’ and ‘cds’. The version used for comparison is noted below in each evaluation step. To our knowledge, the NCGR took extra post-processing steps to filter content, leaving only coding sequences in the ‘cds’ versions of each assembly [31]

### 2. Perform quality control

Reads were analyzed with FastQC (version 0.11.5) [44] and multiqc (version 1.2) [45] to confirm overall qualities before and after trimming. A conservative trimming approach [46] was used with Trimmomatic (version 0.33) [28] to remove residual Illumina adapters and cut bases off the start (LEADING) and end (TRAILING) of reads if they were below a threshold Phred quality score (Q<2).

### 3. Apply digital normalization

To decrease the memory requirements for each assembly, digital normalization was applied with the khmer software package (version 2.0) prior to assembly [47]. First, reads were interleaved, normalized to a k-mer (k = 20) coverage of 20 and a memory size of 4e9, then low-abundance k-mers from reads with a coverage above 18 were trimmed. Orphaned reads, where the mated pair was removed during normalization, were added to the normalized reads.

### 4. Assemble

Transcriptomes were assembled from normalized reads with Trinity 2.2.0 using default parameters (k = 25). This version of Trinity (2.2.0) did not include the “in silico normalization” option as a default parameter. The digital normalization approach we used with khmer is the same algorithm implemented in Trinity, but it requires less memory and is faster [48].

The resulting assemblies are referred to below as the “Lab for Data Intensive Biology” assemblies, or DIB assemblies. The original assemblies are referred to as the NCGR assemblies.

### 5. Post-assembly assessment

Transcriptomes were annotated using the dammit pipeline [49], which relies on the following databases as evidence: Pfam-A (version 28.0) [50], Rfam (version 12.1) [51], OrthoDB (version 8) [52]. In the case where there were multiple database hits, one gene name per contig was selected by choosing the name of the lowest e-value match (<1e-05).

All assemblies were evaluated using metrics generated by the Transrate program [53]. Trimmed reads were used to calculate a Transrate score for each assembly, which represents the geometric mean of all contig scores multiplied by the proportion of input reads providing positive support for the assembly [50]. Comparative metrics were calculated using Transrate for each MMETSP sample between DIB and the NCGR assemblies using the Conditional Reciprocal Best BLAST hits (CRBB) algorithm [54]. A forward comparison was made with the NCGR assembly used as the reference and each DIB re-assembly as the query. Reverse comparative metrics were calculated with each DIB re-assembly as the reference and the NCGR assembly as the query. Transrate scores were calculated for each assembly using the Trimmomatic quality-trimmed reads, prior to digital normalization.

Benchmarking Universal Single-Copy Orthologs (BUSCO) software (version 3) was used with a database of 215 orthologous genes specific to protistans and 303 genes specific to eukaryota with open reading frames in the assemblies. BUSCO scores are frequently used as one measure of assembly completeness [55]

To assess the occurrences of fixed-length words in the assemblies, unique 25-mers were measured in each assembly using the HyperLogLog (HLL) estimator of cardinality built into the khmer software package [56]. We used the HLL function to digest each assembly and count the number of distinct fixedlength substrings of DNA (k-mers).

Unique gene names were compared from a random subset of 296 samples using the dammit annotation pipeline [49]. If a gene name was annotated in NCGR but not in DIB, this was considered a gene uniquely annotated in NCGR. Unique gene names were normalized to the total number of annotated genes in each assembly.

A Tukey’s honest significant different (HSD) post-hoc range test of multiple pairwise comparisons was used in conjunction with an ANOVA to measure differences between distributions of data from the top eight most-represented phyla (“Bacillariophyta”, “Dinophyta”, “Ochrophyta”, “Haptophyta”, “Ciliophora”, “Chlorophyta”, “Cryptophyta”, “Others”) using the ‘agricolae’ package version 1.2-8 in R version 3.4.2 (2017-09-28). Margins sharing a letter in the group label are not significantly different at the 5% level (8). Averages are reported ± standard deviation.

## Results

After assemblies and annotations were completed, files were uploaded to Figshare and Zenodo are available for download [57]. Due to obstacles encountered uploading and maintaining 678 assemblies on Figshare, Zenodo will be the long-term archive for these re-assemblies http://doi.org/10.5281/zenodo.1212585. Assembly quality metrics were summarized and are available (Supplemental Tables 1 and 2 [43]).

**Table 1.**
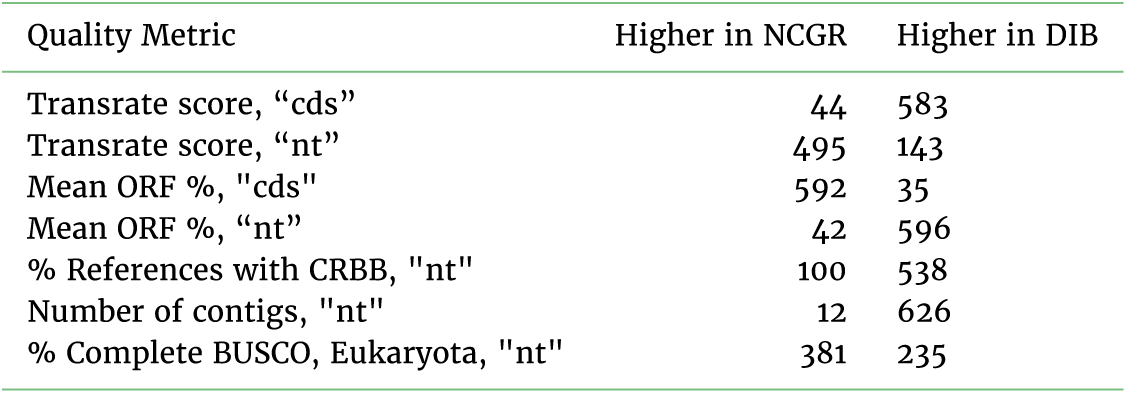
Number of assemblies with higher values in NCGR or DIB for each quality metric. The “cds” or “nt” indicate the version of the NCGR assembly compared with. The NCGR “cds” assemblies were filtered for ORF content.

### Diferences in available evaluation metrics between NCGR and DIB were variable

The majority of transcriptome evaluation metrics collected for each sample were higher in Trinity-based DIB re-assemblies than for the Trans-ABySS-based NCGR assemblies, ‘cds’ versions (1). The Transrate score from the “nt” version of the assemblies were higher in NCGR vs. DIB, whereas compared to the ‘cds’ version, the DIB re-assemblies were higher (Supplemental Figure 1 [43]). Since the NCGR ‘cds’ assemblies were filtered for open reading frame (ORF) content, and the DIB reassemblies were not filtered, the unfiltered NCGR ‘nt’ assemblies are more comparable to the DIB re-assemblies.

The DIB re-assemblies had more contigs than the NCGR assemblies in 83.5% of the samples (1). The mean number of contigs in the DIB re-assemblies was 48,361 ± 35,703 while the mean number of contigs in the NCGR ‘nt’ assemblies was 30,532 ± 21,353 (2). A two-sample Kolmogorov-Smirnov test comparing distributions indicated that the number of contigs were significantly different between DIB and NCGR assemblies (p < 0.001, D = 0.35715). Transrate scores [53], which calculate the overall quality of the assembly based on the original reads, were significantly higher in the DIB re-assemblies (0.31 ± 0.1) compared to the ‘cds’ versions of the NCGR assemblies (0.22 ± 0.09) (p < 0.001, D = 0.49899). The Transrate scores in the NCGR ‘nt’ assemblies (0.35 ± 0.09) were significantly higher than the DIB assemblies (0.22 ± 0.09) (p < 0.001, D = 0.22475) (Supplemental Figure 1 [43]). The frequency of the differences between Transrate scores in the NCGR ‘nt’ assemblies and the DIB re-assemblies is centered around zero (Figure 2C). Transrate scores from the DIB assemblies relative to the NCGR ‘nt’ assemblies did not appear to have taxonomic trends (Supplemental Figure 2 [43]

**Figure 2.**
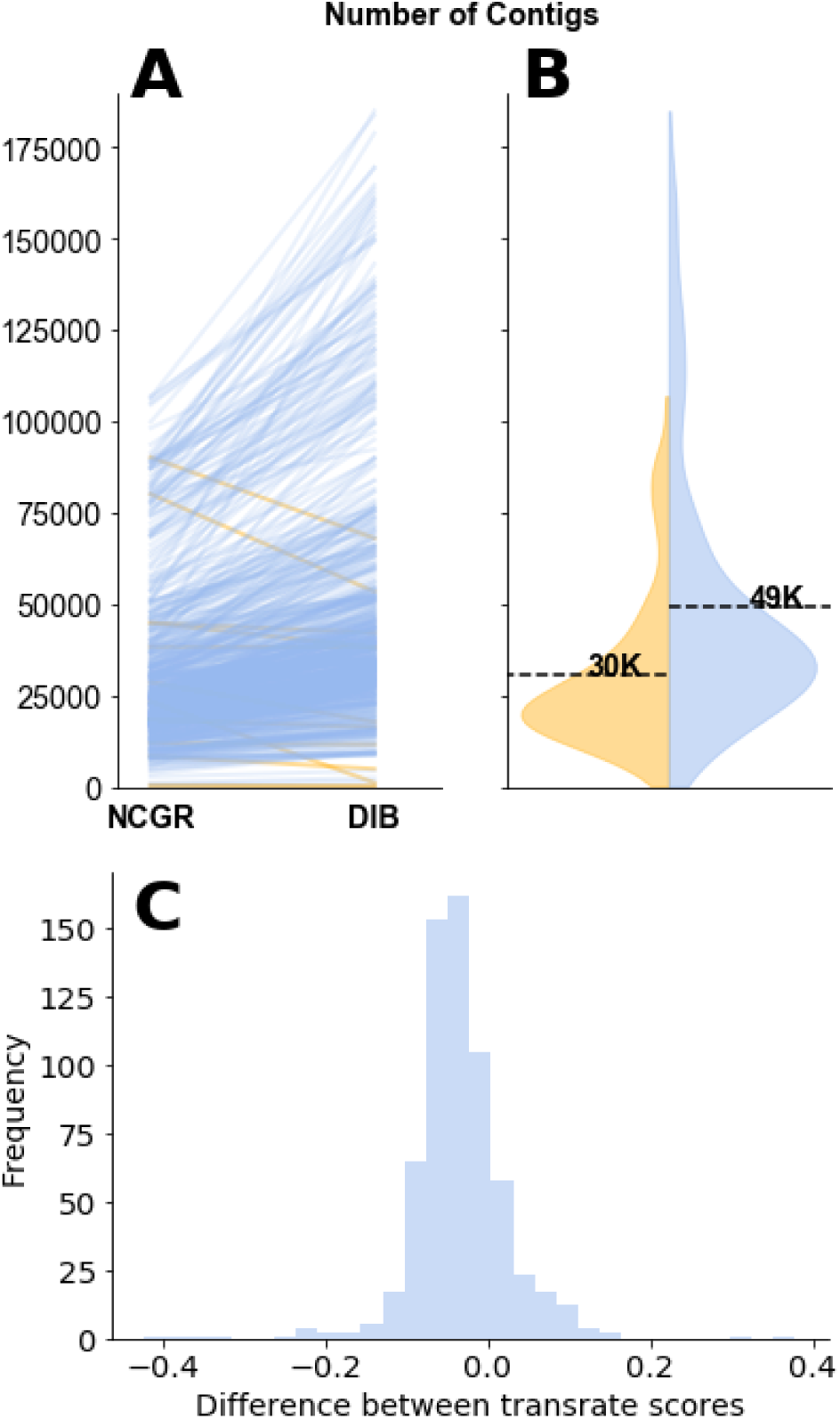
The number of contigs and Transrate quality score for each data set varied between DIB and NCGR assemblies. (A) Slopegraphs show shifts in the number of contigs for each individual sample between the DIB and the NCGR assembly pipelines. Negative slope (yellow) lines represent values where NCGR was higher than DIB and positive slope (blue) lines represent values where DIB was higher than NCGR. (B) Split violin plots show the distribution of the number of contigs in each assembly with the original assemblies from NCGR in yellow (left) and the DIB re-assemblies and in blue (right side of B). (C) The difference in Transrate score between the DIB and NCGR assemblies is shown as a histogram. Negative values on the x-axis indicate that the NCGR assembly had a higher Transrate score and positive values indicate that the DIB assembly had a higher Transrate score.

### The DIB re-assemblies contained most of the NCGR contigs as well as new content

We applied CRBB to evaluate overlap between the assemblies. A positive CRBB result indicates that one assembly contains the same contig information as the other. Thus, the proportion of positive CRBB hits can be used as a scoring metric to compare the relative similarity of content between two assemblies. For example, MMETSP0949 (*Chattonella subsalsa*) had 39,051 contigs and a CRBB score of 0.71 in the DIB re-assembly whereas in the NCGR assembly of the same sample had 18,873 contigs and a CRBB score of 0.34. This indicated that 71% of the reference of DIB was covered by the NCGR assembly, whereas in the reverse alignment, the NCGR reference assembly was only covered by 34% of the DIB re-assembly. The mean CRBB score in DIB when queried against NCGR ‘nt’ as a reference was 0.70 ± 0.22, while the mean proportion for NCGR ‘nt’ assemblies queried against DIB re-assemblies was 0.49 ± 0.10 (p < 0.001, D = 0.71121) (3). This indicates that more content from the NCGR assemblies was included in the DIB re-assemblies than vice versa and also suggests that the DIB re-assemblies overall have additional content. This finding is reinforced by higher unique k-mer content found in the DIB re-assemblies compared to NCGR, where more than 95% of the samples had more unique k-mers in the DIB re-assemblies compared to NCGR assemblies (4).

To investigate whether the new sequence content was genuine, we examined two different metrics that take into account the biological quality of the assemblies. First, the estimated content of open reading frames (ORFs), or coding regions, across contigs was quantified. Though DIB re-assemblies had more contigs, the ORF content is similar to the original assemblies, with a mean of 81.8 ± 9.9% ORF content in DIB re-assemblies and 76.7 ± 10.1% ORF content in the NCGR assemblies. Nonetheless, ORF content in DIB re-assemblies was higher than NCGR assemblies for 95% of the samples (5), although DIB re-assemblies had significantly higher ORF content (p < 0.001, D = 2681). Second, when the assemblies were queried against the eukaryotic BUSCO database [55], the percentages of BUSCO eukaryotic matches in the DIB reassemblies (61.8 ± 19.9%) were similar to the original NCGR assemblies (63.8 ± 20.3%) (5). However, the DIB re-assemblies were significantly different compared to the NCGR assemblies (p = 0.002408, D = 0.099645). Therefore, although the number of contigs and amount of CRBB content were dramatically increased in the DIB re-assemblies compared to the NCGR assemblies, the differences in ORF content and BUSCO matches compared to eukaryotic (5) and protistan (Supplemental Figure 3 [43]) databases, while they were significantly different, were less dramatic. This suggests that content was not lost by gaining extra contigs. Since the extra content contained roughly similar proportions of ORFs and BUSCO annotations, it is likely that the re-assemblies contribute more biologically meaningful information.

**Figure 3.**
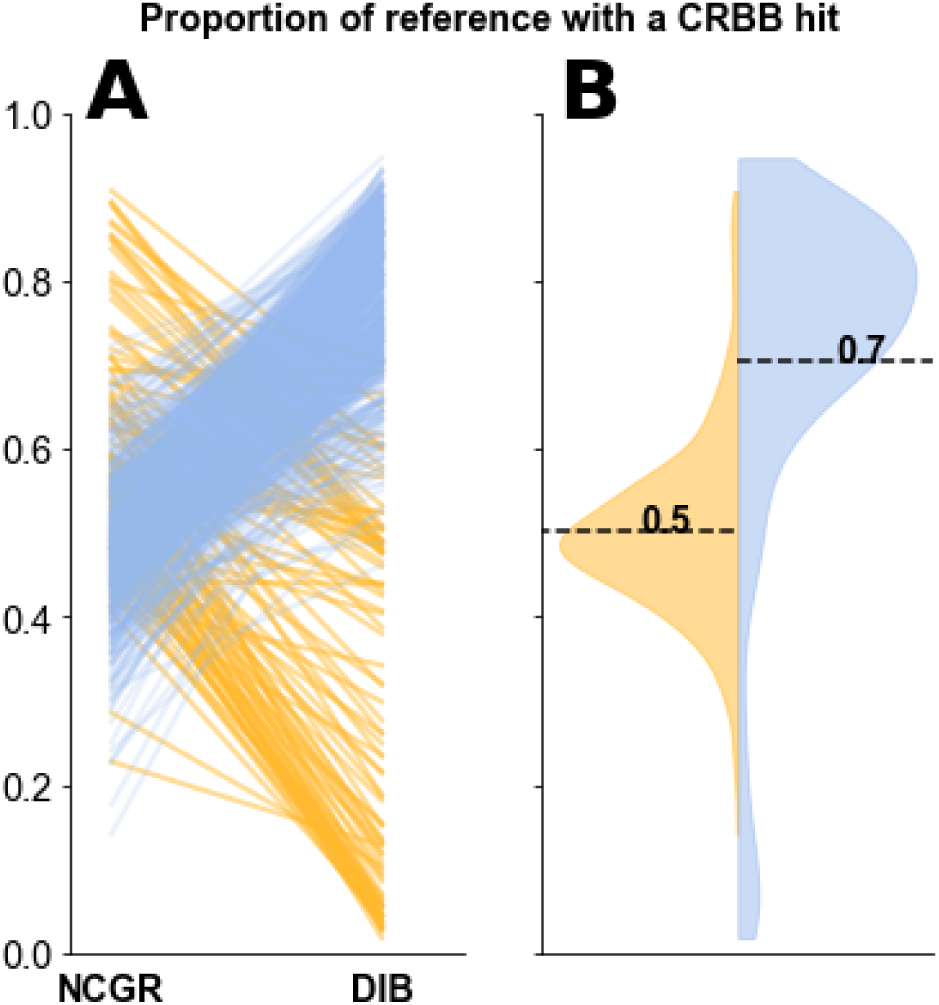
(A) Slopegraphs comparing the proportion of CRBB hits between NCGR ‘nt’ assemblies and DIB assemblies between the same samples. (B) Violin plots showing the distribution of the proportion of NCGR transcripts with reciprocal BLAST hits to DIB (blue) and the proportion of DIB transcripts with reciprocal BLAST hits to NCGR (yellow).

Looking through the results for missing BUSCOs in the eight samples where NCGR had >30% higher complete BUSCO evaluation score (MMETSP0121, MMETSP0932, MMETSP0045, MMETSP0169, MMETSP0232, MMETSP0439, MMETSP0329, MMETSP0717), in some cases a particular orthogroup in the BUSCO database does not produce output for reasons that we don’t understand. For example, the Trinity-based pipeline only produced 342 contigs for sample MMETSP0232 while the NCGR ‘nt’ assembly had 4234 and the ‘cds’ version had 2736. BUSCO did not recognize any of the DIB contigs but it did recognize the NCGR contigs. For other samples, MMETSP0169 (*Corethron pennatum*, Phylum: Bacillariophyta), the BUSCO software recognized several DIB contigs but the BUSCO group was still considered “missing”, even though there were lengths of the contig identified in the output as being similar. For example, the BUSCO orthogroup “EOG0937060I” is a “DNARNA helicase, ATP-dependent, DEAH-box type, conserved site”. The BUSCO output indicates the DIB contig, “TRINITY_DN13758_c3_g2_i1” with a length 974 bases is related to this orthogroup. When we look for this gene in the gff annotation file for MMETSP0169, there are no annotation results for this contig. Another DIB contig, “TRINITY_DN3716_c0_g1_i1” (length 154) is also identified as similar to this same orthogroup. This contig does have annotation results, but it matches with a BUSCO orthogroup, “EOG090C08EI”, which is a different gene, Abl-interactor, homeo-domain homologous domain (ABI family, member 3a). The top results comparing the contig sequence, “TRINITY_DN13758_c3_g2_i1” against the NCBI blastn database matches with small, several hundred bp regions of the EOG0937060I gene sequence (XM_021257656.1, XM_004843976.2, XM_010604294.2, XM_010604293.2, XM_010604291.2, XR_776390.2). Even though this contig was assembled, it did not successfully annotate. We do not know whether there are errors associated with this assembled contig, or if the contig sequence is unique to this MMETSP0169 organism. Since the BUSCO database and corresponding orthogroups were constructed from multiple sequence alignments with existing individuals in the databases, it is possible that the transcriptome from the newly sequenced, MMETSP0169 (*Corethron pennatum*) may naturally fall outside the hmm scoring cutoffs for matching with the BUSCO orthogroups. Since the corresponding NCGR assembly had a “Duplicated” result from this particular BUSCO, it is also possible that there is a particular oddity within this ortholog.

There are many examples that can be picked over in these results, which suggests that there is more to learn about the evaluation tools within the context of the organisms in this data set. For now, we conclude that our assemblies are differently fragmented in some regions relative to the NCGR assemblies. We have assembled additional sequences that were not assembled by NCGR. Some NCGR assemblies had different and more complete content than the DIB assemblies. As far as we can tell, there does not appear to be a pattern in the samples that fared well with this pipeline vs. NCGR. This could be a future avenue to explore.

Following annotation by the dammit pipeline [49], 91 ± 1.6% of the contigs in the DIB re-assemblies had positive matches with sequence content in the databases queried (Pfam, Rfam, and OrthoDB), with 48 ± 0.9% of those containing unique gene names (the remaining are fragments of the same gene). Of those annotations, 7.8 ± 0.2% were identified as novel compared to the NCGR ‘nt’ assemblies, determined by a “false” CRBB result (6). Additionally, the number of unique gene names in DIB re-assemblies were higher in 97% of the samples compared to NCGR assemblies, suggesting an increase in genic content (7).

Novel contigs in the DIB re-assemblies likely represent a combination of unique annotations, allelic variants and alternatively spliced isoforms. For example, “F0XV46_GROCL”, “Helicase_C”, “ODR4-like”,“PsaA_PsaB”, and “Metazoa_SRP” are novel gene names found annotated in the DIB re-assembly of the sample MMETSP1473 (Stichococcus sp.) that were absent in the NCGR assembly of this same sample. Other gene names, for example “Pkinase_Tyr”,“Bromodomain”, and “DnaJ”, are found in both the NCGR and DIB assemblies, but are identified as novel contigs based on negative CRBB results in the DIB re-assembly of sample MMETSP1473 compared to the NCGR reference.

### Assembly metrics varied by taxonomic group being assembled

To examine systematic taxonomic differences in the assemblies, metrics for content and assembly quality were assessed (8). Metrics were grouped by the top eight most represented phyla in the MMETSP data set as follows: Bacillariophyta (N=173), Dinophyta (N=114), Ochrophyta (N=73), Chlorophyta (N=62), Haptophyta (N=61), Ciliophora (N=25), Cryptophyta (N=22) and Others (N=130).

While there were no major differences between the phyla in the number of input reads (Figure 8 A), the Dinoflagellates (Dinophyta) had significantly different (higher) number of contigs (p < 0.01), unique k-mers (p < 0.001), and % ORF (p < 0.001) compared to other groups (8), and assemblies from Ciliates (Ciliophora) had lower % ORF (p < 0.001) (8).

## Discussion

### DIB re-assemblies contained the majority of the previously-assembled contigs

We used a different pipeline than the original one used to create the NCGR assemblies, in part because new software was available [18] and in part because of new trimming guidelines [46]. The general genome assembler ABySS [20] was used in conjunction with a *de novo* transcriptome assembly pipeline described by Keeling et al. [31]. We had no a priori expectation for the similarity of the results, yet we found that the majority of new DIB re-assemblies included substantial portions of the previous NCGR assemblies seen in the CRBB results. Given this, it may seem surprising that the Transrate and BUSCO scores are lower in the DIB re-assemblies relative to the NCGR counterparts. However, given that the number of contigs and the k-mer content were both dramatically increased in the DIB reassemblies, it is interesting that the ORFs and annotations were similar between the two assemblies. If the extra content observed was due to assembly artifact, we would not expect these content-based results to be similar. The two metrics, Transrate and BUSCO, which estimate “completeness” of the transcriptomes, may not be telling the whole story. Our results suggest that both pipelines yielded similarly valid contigs, even though the NCGR assemblies appeared to be less sensitive.

The relative increase in number of unique k-mers from the NCGR assemblies to the DIB re-assemblies could be due to the higher number of contigs generated by Trinity. Within the data, the Trinity assembler found evidence for building alternative isoforms. The ABySS assembler and transcriptome pipeline that NCGR used [31] appears to not have preserved that variation, perhaps in an attempt to narrow down the contigs to a consensus transcript sequence.

### Re-assembly with new tools can yield new results

Evaluation with quality metrics suggested that the DIB reassemblies were more inclusive than the NCGR assemblies. The Transrate scores in the DIB re-assemblies compared to the NCGR ‘nt’ assemblies were significantly lower, indicating that the NCGR ‘nt’ assemblies had better overall read inclusion in the assembled contigs whereas the DIB assemblies had higher Transrate scores than the NCGR ‘cds’ version. This suggests that the NCGR ‘cds’ version, which was post-processed to only include coding sequence content, was missing information originally in the quality-trimmed reads. As we also saw with % ORF, when filtration steps select only for ORF content in the NCGR ‘cds’ versions, potentially useful content is lost. The Transrate score [53] is one of the few metrics available for evaluating the ‘quality’ of a *de novo* transcriptome. It is similar to the DETONATE RSEM-EVAL score in that it returns a metric indicating how well the assembly is supported by the read data [13]. It does not directly evaluate the underlying de Bruijn graph data structure used to produce the assembled contigs. In the future, metrics directly evaluating the underlying de Bruijn graph data structure may better evaluate assembly quality. Here, the DIB re-assemblies, which used the Trinity *de novo* assembly software, typically contained more k-mers, more annotated transcripts, and more unique gene names than the NCGR assemblies.

These points all suggest that additional content in these re-assemblies might be biologically relevant and that these reassemblies provide new content not available in the previous NCGR assemblies. Since contigs are probabilistic predictions of full-length transcripts made by assembly software [18], ‘final’ reference assemblies are approximations of the full set of transcripts in the transcriptome. Results from this study suggest that the ‘ideal’ reference transcriptome is a moving target and that these predictions may continue to improve given updated tools in the future.

For some samples, complete BUSCO scores were lower than over half of DIB vs. NCGR. This could be an effect of the BUSCO metric, given that these samples did not perform poorly with other metrics such as % ORF and number of contigs compared to the NCGR. For other samples, MMETSP0252 (*Prorocentrum lima*) in particular, assemblies required several tries and only four contigs were assembled from 30 million reads. The fastqc reports were unremarkable, compared to the other samples. In such a large dataset with a diversity of species with no prior sequencing data it is challenging to speculate why each anomaly occurred. However, further investigation into the reasons for failures and peculiarities in the evaluation metrics may lead to interesting discoveries about how we should be effectively assembling and evaluating nucleic acid sequencing data from a diversity of species.

We predict that assembly metrics could have been further improved with longer read lengths of the original data since MMETSP data had only 50 bp read lengths, although this would have presented Keeling et al. [31] with a more expensive data collection endeavor. A study by Chang et al. [25] reported a consistent increase in the percentage of full-length transcript reconstruction and a decrease in the false positive rate moving from 50 to 100 bp read lengths with the Trinity assembler. However, regardless of length, the conclusions we draw would likely remain the same that assembling data with new tools can yield new results.

The DIB re-assemblies, including the additional biologically relevant information, are likely to be meta-transcriptomes. RNA sequences generated from the MMETSP experiments are likely to contain genetic information from more than the target species, as many were not or could not be cultured axenically. Thus, both the NCGR assemblies and the DIB reassemblies, including the additional biologically relevant information, might be considered meta-transcriptomes. Sequencing data and unique k-mer content likely include bacteria, viruses, or other protists that occurred within the sequenced sample. We did not make an attempt to de-contaminate the assemblies.

The evaluation metrics described here generally serve as a framework for better contextualizing the quality of protistan transcriptomes. For some species and strains in the MMETSP data set, these data represent the first nucleic acid sequence information available [31].

### Automated and programmable pipelines can be used to process arbitrarily many RNAseq samples

The automated and programmable nature of this pipeline was useful for processing large data sets like the MMETSP as it allowed for batch processing of the entire collection, including reanalysis when new tools or new samples become available (see op-ed Alexander et al. 2018). During the course of this project, we ran two re-assemblies of the complete MMETSP data set and one subset as new versions Trinity were released (Supplemental Notebook 2 [43]). Each re-analysis of the complete dataset required only a single command and approximately half a CPU-year of compute. The value of automation is clear when new data from samples become available to expand the data set, tools are updated, or many tools are compared in benchmark studies. Despite this, few assembly efforts completely automate their process, perhaps because the up-front cost of doing so is high compared to the size of the dataset typically being analyzed.

For the purposes of future benchmarking studies, a subset of 12 “High” and 15 “Low” performing samples were identified based on the evaluation metrics: number of contigs, longest contig length, unique k-mers (k=25), and % Complete BUSCO (eukaryota) (Supplemental Figure 4 [43]).

**Figure 4.**
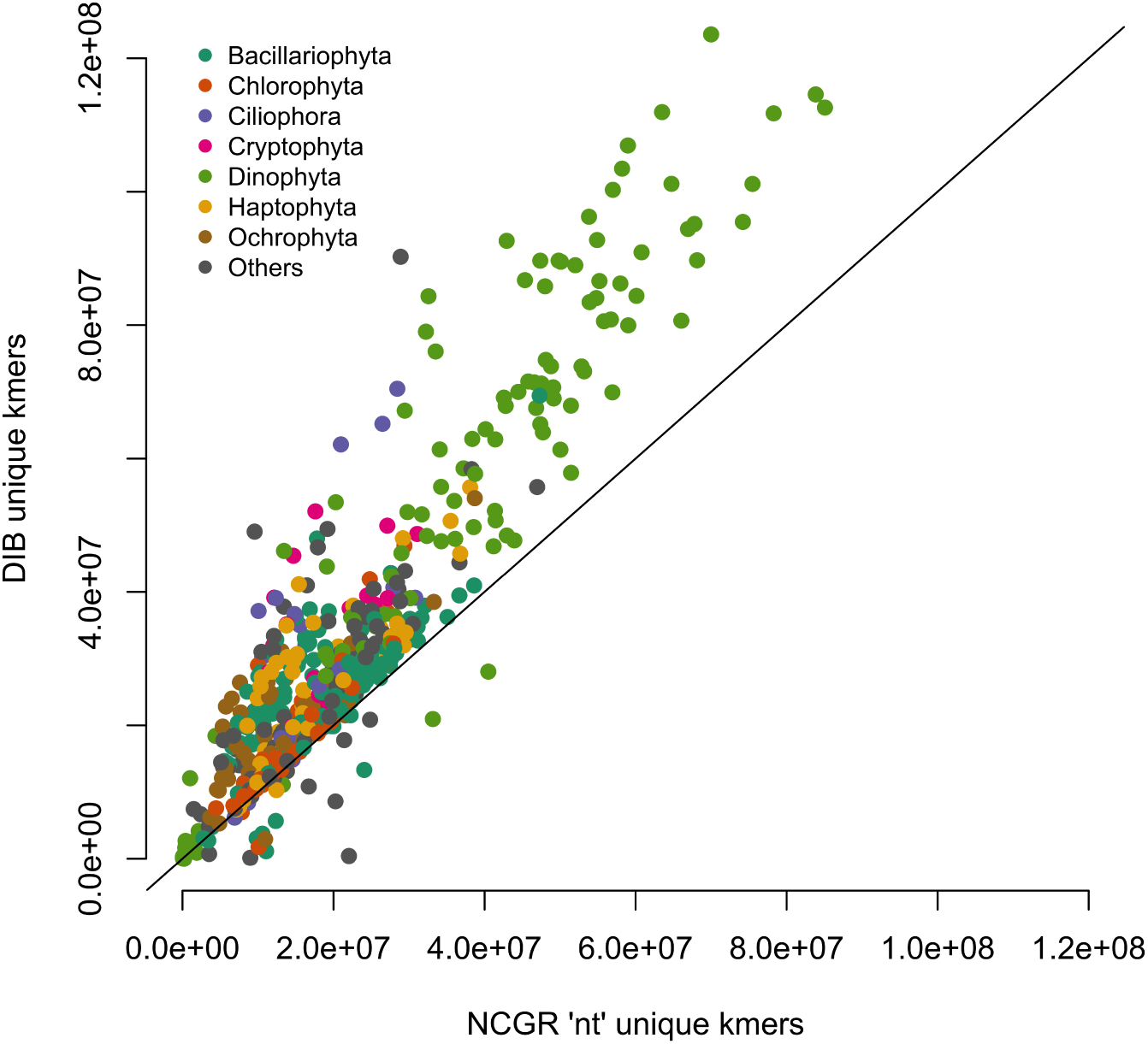
Unique numbers of k-mers (k=25) in seven most represented phyla, calculated with the HyperLogLog function in the khmer software package. DIB re-assemblies were compared to the NCGR ‘nt’ assemblies along a 1:1 line. Samples are colored based on their phylum level afiliation. More than 95% of the DIB re-assemblies had more unique k-mers than to the NCGR assembly of the same sample.

**Figure 5.**
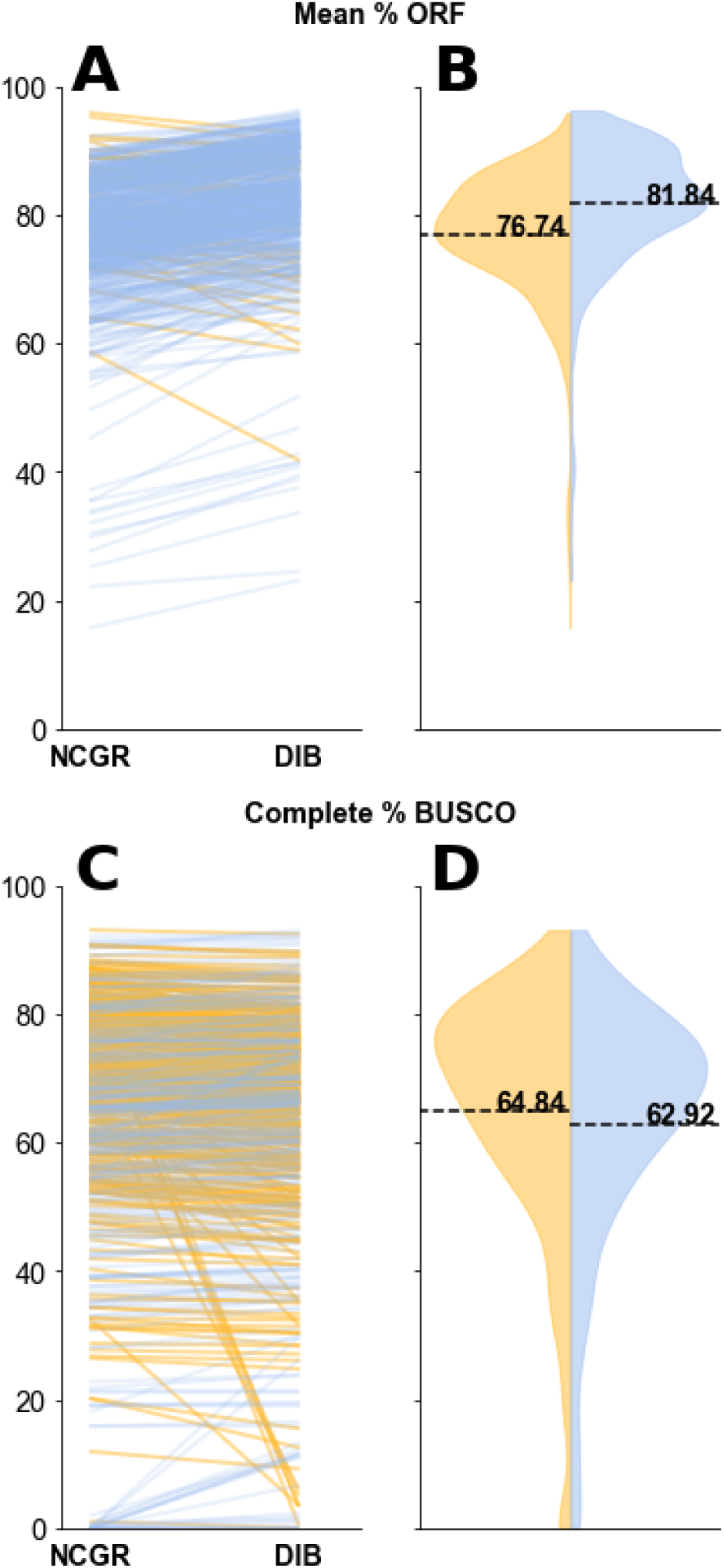
The percentage of contigs with a predicted open reading frame (ORF) (A, B) and the percentage of complete protistan universal single-copy orthologs (BUSCO) recovered in each assembly (C, D). In the blue (right side B, D) are the “DIB” re-assemblies and in yellow (left side of B, D) are the original ‘nt’ assemblies from NCGR. Slopegraphs (A,C) compare values between the DIB and the NCGR ‘nt’ assemblies. Yellow lines represent negative slope values where NCGR was higher than DIB and blue lines represent positive slope values where DIB was higher than NCGR.

**Figure 6.**
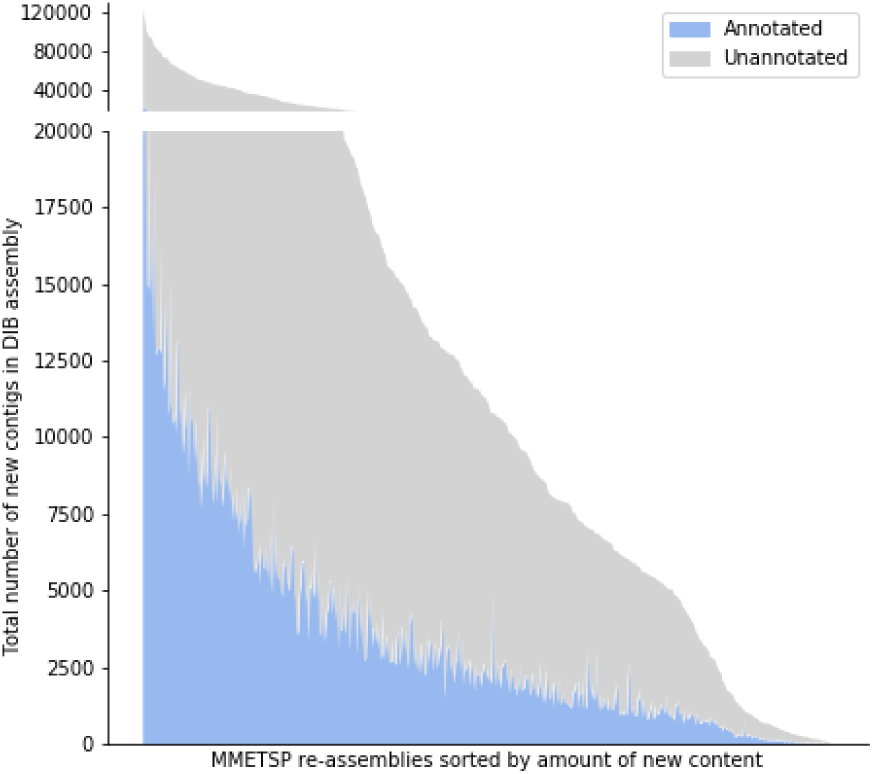
A histogram across MMETSP samples depicting the number of contigs identified as novel in DIB assemblies. These contigs were absent in the NCGR assemblies, based on negative conditional reciprocal best BLAST (CRBB) results. Samples are sorted from highest to lowest number of ‘new’ contigs. The region in gray indicates the number of unannotated contigs present in the DIB re-assemblies, absent from NCGR ‘nt’ assemblies. Highlighted in blue are contigs that were annotated with dammit [49] to a gene name in the Pfam, Rfam, or OrthoDB databases, representing the number of contigs unique to the DIB re-assemblies with an annotation.

**Figure 7.**
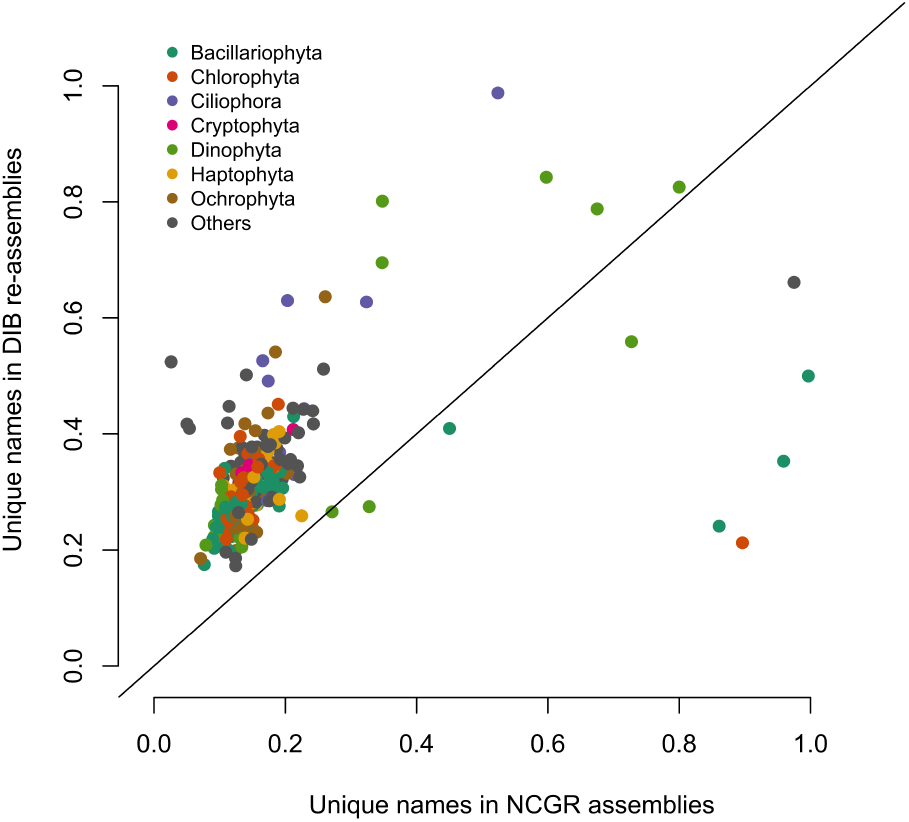
Unique gene names found in a subset (296 samples) of either NCGR ‘nt’ assemblies or DIB re-assemblies but not found in the other assembly, normalized to the number of annotated contigs in each assembly. The line indicates a 1:1 relationship between the unique gene names in DIB and NCGR. More than 97% of the DIB assemblies had more unique gene names than in NCGR assemblies of the same sample.

**Figure 8.**
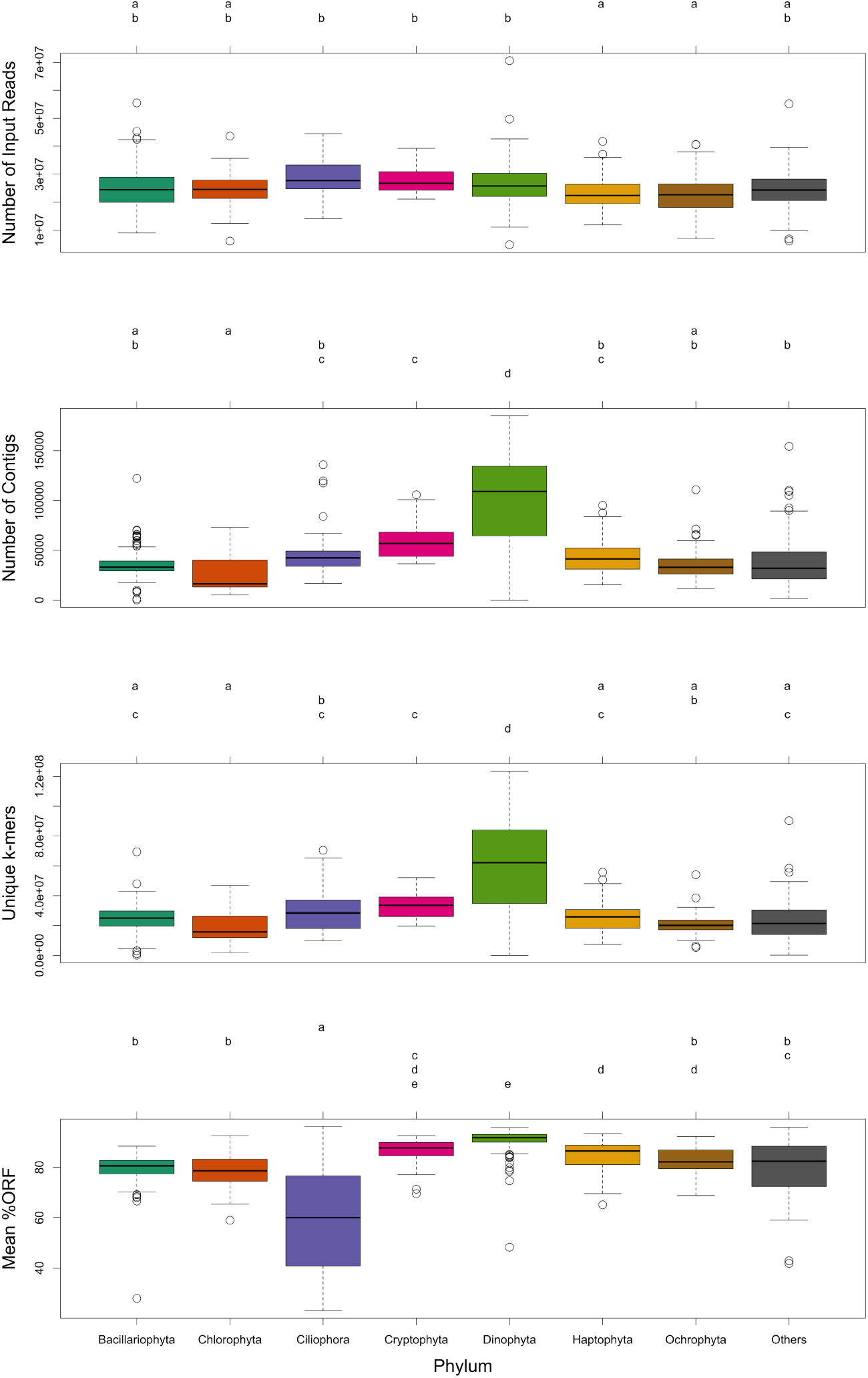
Box-and-whisker plots for the seven most common phyla in the MMETSP dataset, (A) number of input reads, (B) number of contigs in the assembly, (C) unique k-mers (k = 25) in the assembly, (D) mean percentage open reading frames (ORF). Groups sharing a letter in the top margin were generated from Tukey’s HSD post-hoc range test of multiple pairwise comparisons used in conjunction with an ANOVA.

### Analyzing many samples using a common pipeline identfies taxonspecfic trends

The MMETSP dataset presents an opportunity to examine transcriptome qualities for hundreds of taxonomically diverse species spanning a wide array of protistan lineages. This is among the largest set of diverse RNAseq data to be sequenced. In comparison, the Assemblathon2 project compared genome assembly pipelines using data from three vertebrate species [58]. The BUSCO paper assessed 70 genomes and 96 transcriptomes representing groups of diverse species (vertebrates, arthropods, other metazoans, fungi) [55]. Other benchmarking studies have examined transcriptome qualities for samples rep-resenting dozens of species from different taxonomic groupings [59, 60]. A study with a more restricted evolutionary analysis of 15 plant and animals species [60] found no evidence of taxonomic trends in assembly quality but did find evidence of differences between assembly software packages [59].

With the MMETSP data set, we show that comparison of assembly evaluation metrics across this diversity provides not only a baseline for assembly performance, but also highlights particular metrics which are unique within some taxonomic groups. For example, the phyla Ciliophora had a significantly lower percentage of ORFs compared to other phyla. This is supported by recent work which has found that ciliates have an alternative triplet codon dictionary, with codons normally encoding STOP serving a different purpose [37, 38, 39], thus application of typical ORF finding tools fail to identify ORFs accurately in Ciliophora. Additionally, Dinophyta data sets had a significantly higher number of unique k-mers and total contigs in assemblies compared to the assemblies from other data sets, despite having the same number of input reads. Such a finding supports previous evidence from studies showing that large gene families are constitutively expressed in Dinophyta [61].

In future development of *de novo* transcriptome assembly software, the incorporation of phylum-specific information may be useful in improving the overall quality of assemblies for different taxa. Phylogenetic trends are important to consider in the assessment of transcriptome quality, given that the assemblies from Dinophyta and Ciliophora are distinguished from other assemblies by some metrics. Applying domain-specific knowledge, such as specialized transcriptional features in a given phyla, in combination with other evaluation metrics can help to evaluate whether a transcriptome is of good quality or “finished” enough to serve as a high quality reference to answer the biological questions of interest.

## Conclusion

As the rate of sequencing data generation continues to increase, efforts to programmatically automate the processing and evaluation of sequence data will become increasingly important. Ultimately, the goal in generating *de novo* transcriptomes is to create the best possible reference against which downstream analyses can be accurately based. This study demonstrated that re-analysis of old data with new tools and methods improved the quality of the reference assembly through an expansion of the gene catalog of the dataset. Notably, these improvements arose without further experimentation or sequencing.

With the growing volume of nucleic acid data in centralized and decentralized repositories, streamlining methods into pipelines will not only enhance the reproducibility of future analyses, but will facilitate inter-comparisons amongst from both similar and diverse datasets. Automation tools were key in successfully processing and analyzing this large collection of 678 samples.

## Acknowledgements

The authors would gratefully like to acknowledge the contributions of Camille Scott (0000-0001-8822-8779), Luiz Irber (0000-0003-4371-9659), Taylor Reiter (0000-0002-7388-421X), Daniel Standage (0000-0003-0342-8531) and other members of the Data Intensive Biology lab at UC Davis for providing their time towards helpful assistance with troubleshooting the assembly, annotation and evaluation pipeline and visualizations presented in this paper.

## Declarations

### List of abbreviations

BLAST: Basic Local Alignment Search Tool
CRBB: Conditional Recriprocal Best BLAST
DIB: Data Intensive Biology Lab at the University of California Davis
HLL: HyperLogLog
MMETSP: Marine Microbial Eukaryotic Transcriptome Sequencing Project
NCGR: National Center for Genome Research
ORF: Open Reading Frame
NCBI: National Center for Biotechnology Information
SRA: Sequence Read Archive

### Ethical Approval

Data were downloaded from public repositories, provided by and NCBI BioProject PRJNA231566 as cited in the text.

### Consent for publication

Not applicable.

### Author’s Contributions

Re-assembly, automated analysis pipeline, and data curation were done by LKJ. Writing, review & editing were done by LKJ, HA, CTB. Visualizations were developed by HA and LKJ.

